## Supplementary Material for "Variability of visual field maps in human early extrastriate cortex challenges the canonical model of organization of V2 and V3"

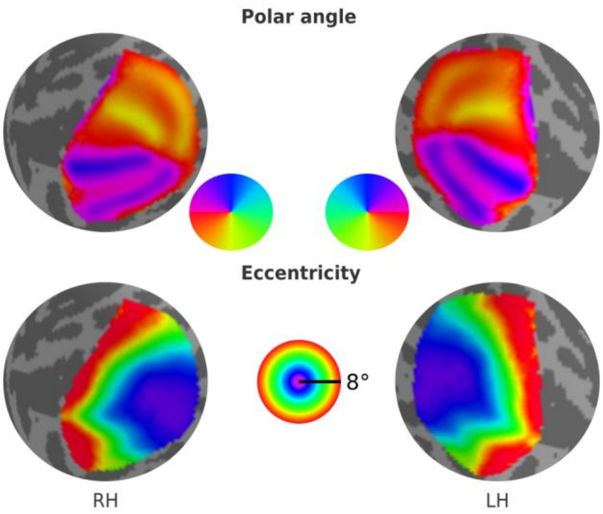


**Supplementary Figure 1 – Average retinotopic maps across all 181 individuals from the HCP retinotopy dataset for both left (LH) and right (RH) hemispheres.**

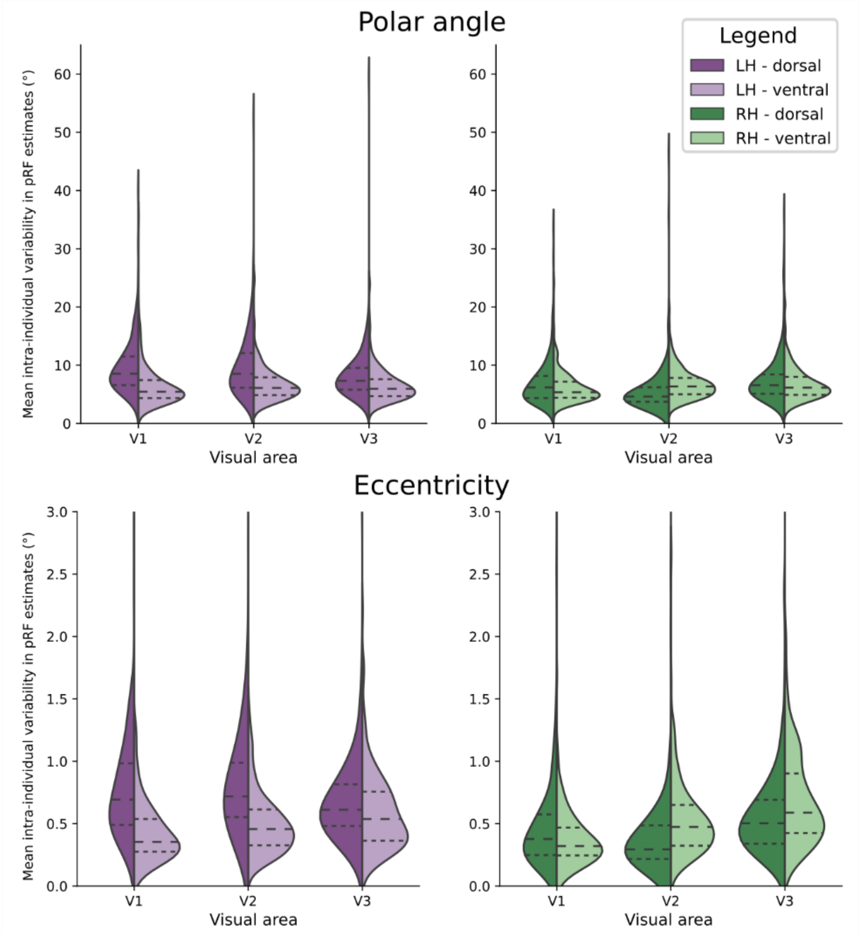


**Supplementary Figure 2 – Intra-individual variability in visual field maps of early visual areas.** Empirical distributions of intra-individual variability of polar angle (top) and eccentricity (bottom) maps for both dorsal (dark shades) and ventral (lighter shades) portions of early visual areas in the left (purple) and right (green) hemispheres. The intra-individual variability is the difference in pRF estimates from two pRF model fits; each used half of the retinotopic mapping data (see Supplementary Table 1 for more information).

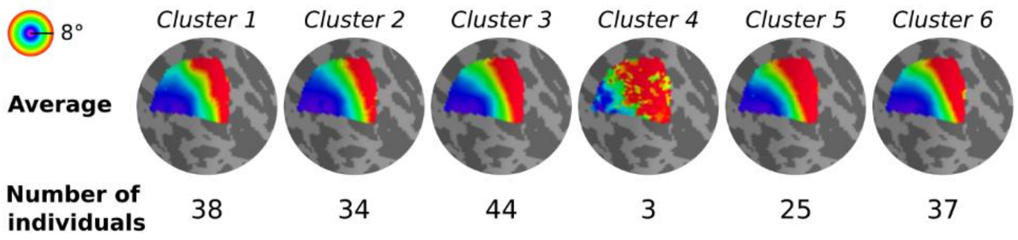


**Supplementary Figure 3 – Clusters of eccentricity maps of the dorsal portion of early visual cortex.** An average continuous map was calculated for each cluster by averaging the continuous eccentricity maps across all individuals within each cluster.

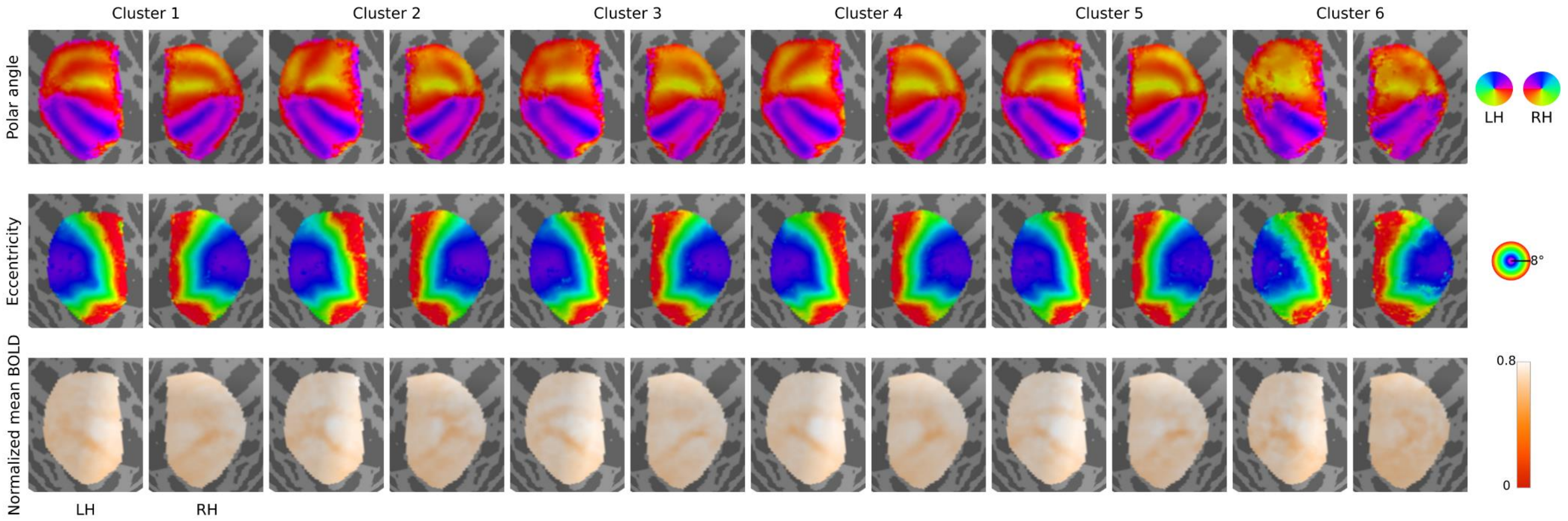


**Supplementary Figure 4 – Average polar angle, eccentricity, and normalized mean BOLD signal maps from each cluster and hemisphere.** Average continuous polar angle, eccentricity, and normalized mean BOLD signal maps were calculated for each cluster by averaging the continuous maps across all individuals within each cluster. As in the manuscript, the cluster assignment was based on the clustering analysis with polar angle maps from the dorsal portion of the early visual cortex and left hemisphere.

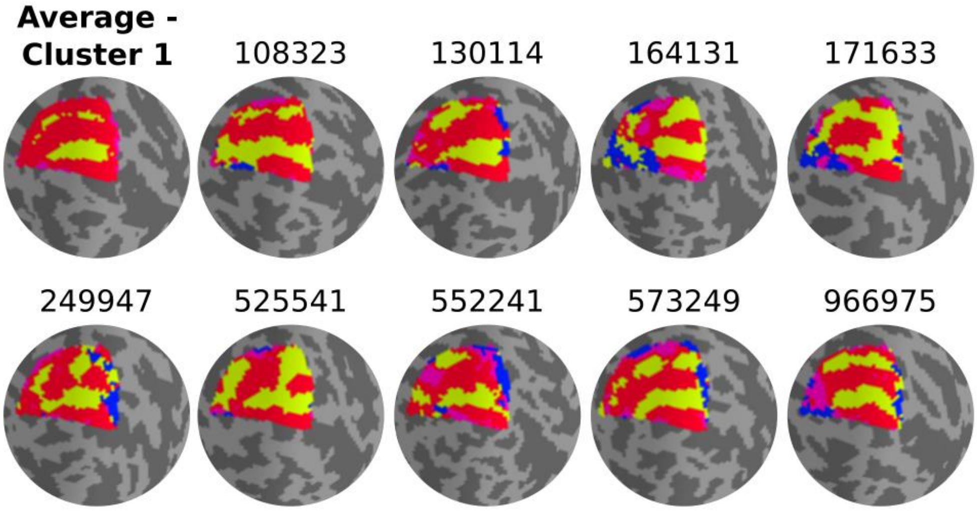


**Supplementary Figure 5 – Nine randomly selected polar angle maps within Cluster 1.** The average discrete map of Cluster 1 is shown.

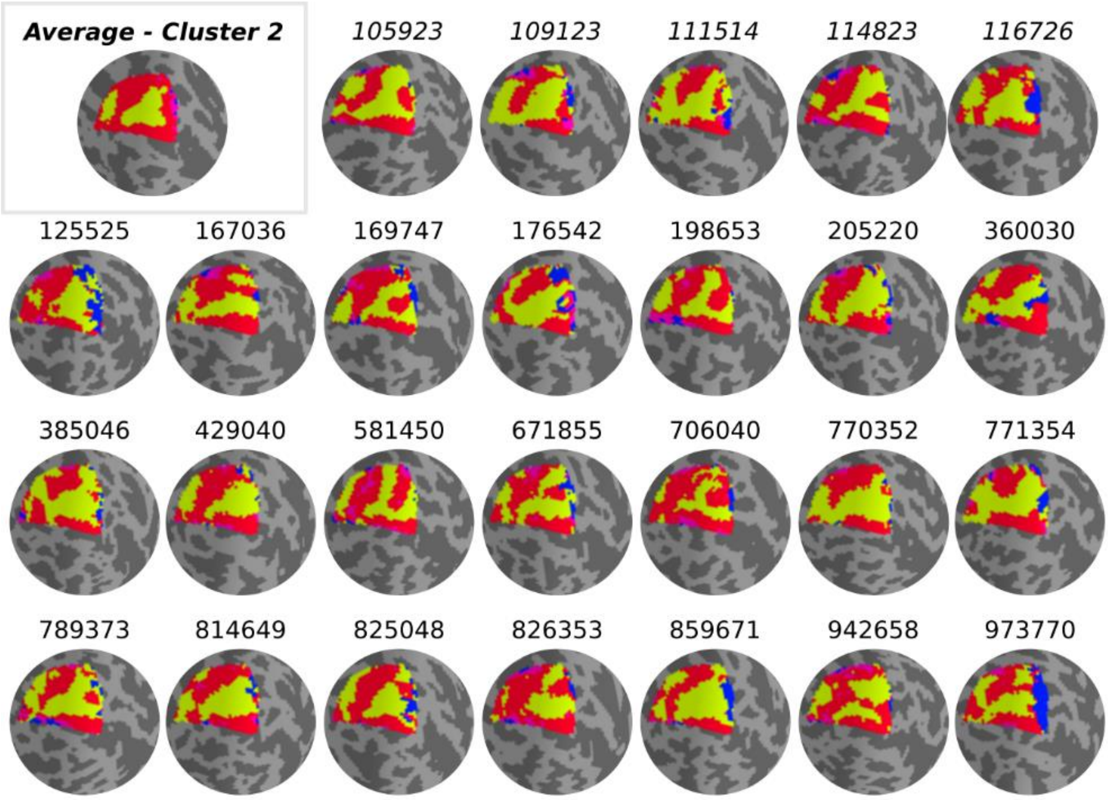


**Supplementary Figure 6 – Individual polar angle maps within Cluster 2.** The average discrete map of Cluster 2 is shown. All individuals' polar angle maps within Cluster 2 (total of 26 individuals) are shown for qualitative examination.

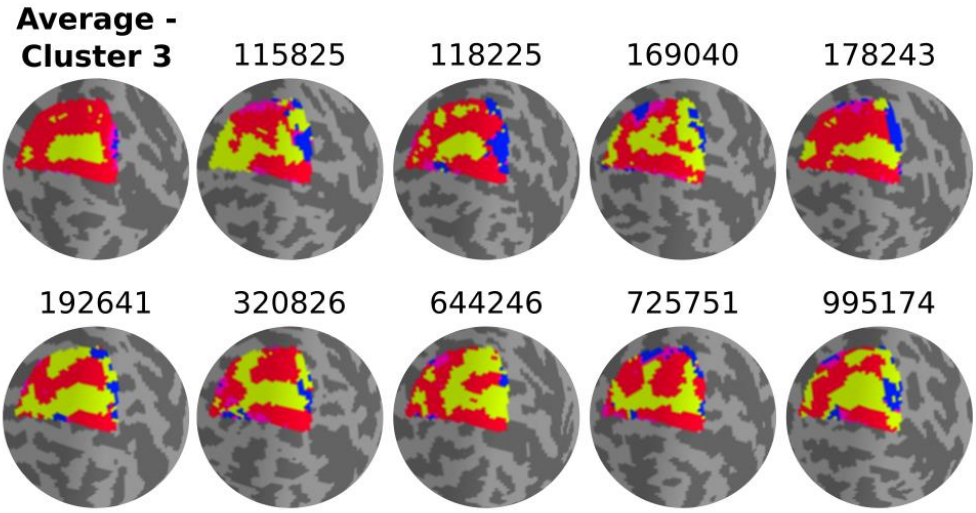


**Supplementary Figure 7 – Nine randomly selected polar angle maps within Cluster 3.** The average discrete map of Cluster 3 is shown.

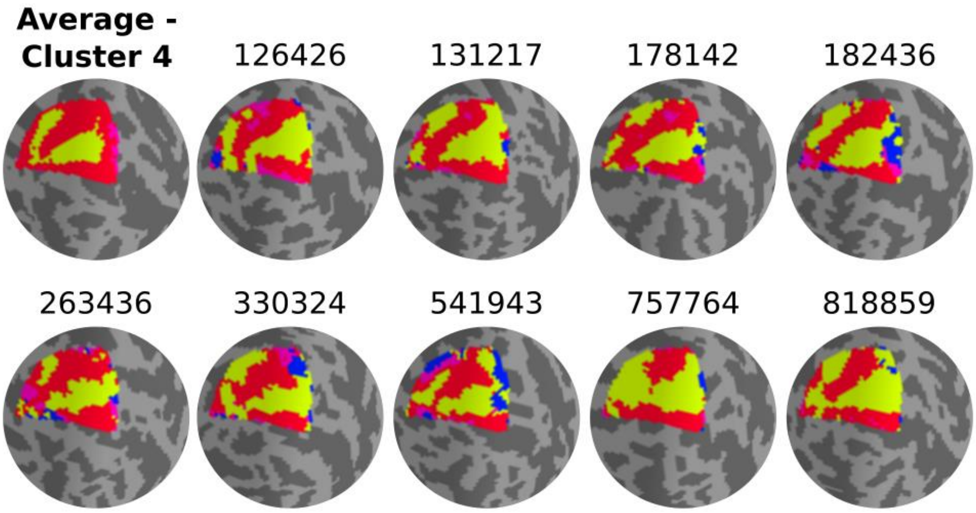


**Supplementary Figure 8 – Nine randomly selected polar angle maps within Cluster 4.** The average discrete map of Cluster 4 is shown.

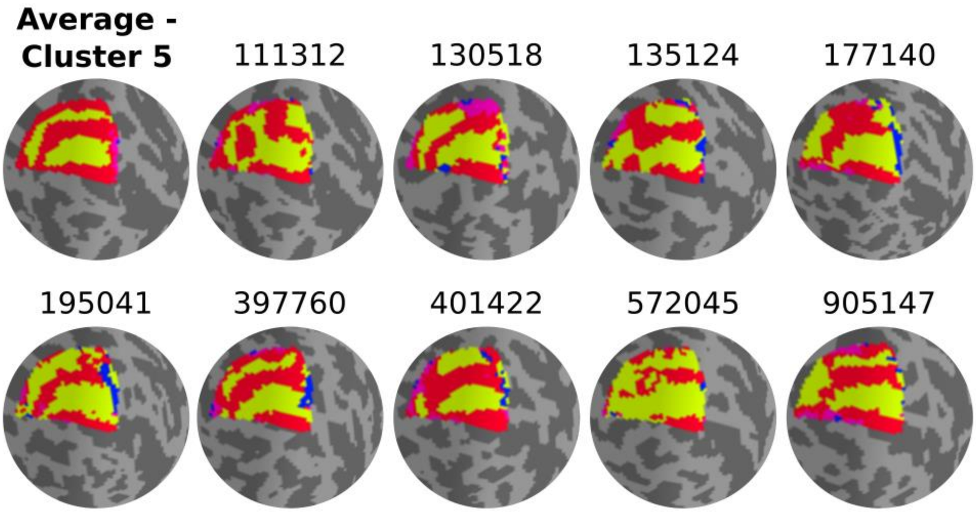


**Supplementary Figure 9 – Nine randomly selected polar angle maps within Cluster 5.** The average discrete map of Cluster 5 is shown.

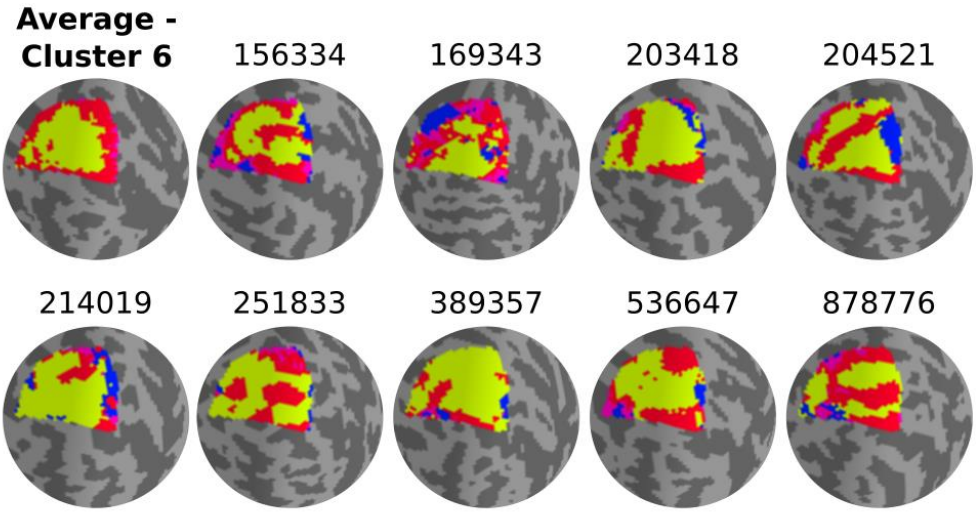


**Supplementary Figure 10 – Nine randomly selected polar angle maps within Cluster 6.** The average discrete map of Cluster 6 is shown.

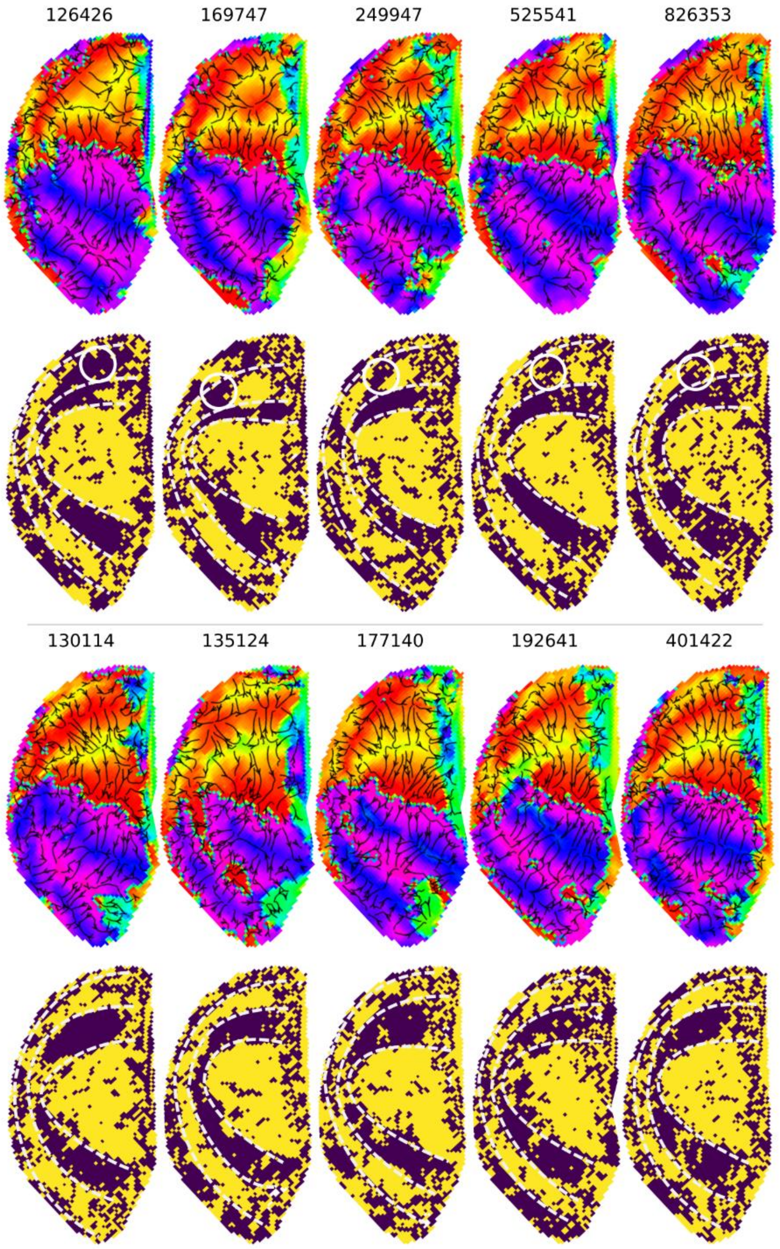


**Supplementary Figure 11 – Visual field sign analysis for delineating visual areas.** Five more examples of left hemisphere polar angle maps with unusual Y-shaped lower vertical representations are shown in the top half. Polar angle gradients are shown in a 'streamline' representation (first row) with their respective visual field sign representation (second row). In the bottom half are five other examples of polar angle maps with a truncated V3 boundary, indicating that dorsal V3 does not cover the entire quarter visual field (i.e., from 360º to 270º). Despite that, the visual field sign representation does not show discontinuity in the typical mirror-image representation of dorsal V3.

**Supplementary Table 1** – Summary of the data used for the analyses described in the main manuscript.

| File | Data type | Analysis |
|---|---|---|
| S1200_7T_Retinotopy181.All.Fit1_PolarAngle_MSMAll.32k_fs_LR.dscalar.nii[1] | Polar angle maps | Individual variability |
| S1200_7T_Retinotopy181.All.Fit1_Eccentricity_MSMAll.32k_fs_LR.dscalar.nii | Eccentricity maps | Individual variability |
| S1200_7T_Retinotopy181.All.curvature_MSMAll.32k_fs_LR.dscalar.nii | Curvature maps | Individual variability / covariate |
| S1200_7T_Retinotopy181.All.Fit1_MeanBOLD_MSMAll.32k_fs_LR.dscalar.nii | Mean BOLD maps | Individual variability / covariate |
| S1200_7T_Retinotopy181.All.**Fit2**_PolarAngle_MSMAll.32k_fs_LR.dscalar.nii | Polar angle maps | Intra-individual variability / covariate |
| S1200_7T_Retinotopy181.All.**Fit3**_PolarAngle_MSMAll.32k_fs_LR.dscalar.nii | Polar angle maps | Intra-individual variability / covariate |
| S1200_7T_Retinotopy181.All.**Fit2**_Eccentricity_MSMAll.32k_fs_LR.dscalar.nii | Eccentricity maps | Intra-individual variability / covariate |
| S1200_7T_Retinotopy181.All.**Fit3**_Eccentricity_MSMAll.32k_fs_LR.dscalar.nii | Eccentricity maps | Intra-individual variability / covariate |

[1] "S1200_7T_Retinotopy181.All.(modality)_MSMAll.32k_fs_LR.dscalar.nii" includes collated data from all 181 participants.

**Supplementary Table 2 – Gaze position change as a function of cluster assignment.** The mean deviation in gaze position along the X and Y axis across runs of retinotopic mapping stimuli presentation and individuals is shown for each cluster. We also show the number of individuals with eye-tracking data per cluster, given that eye-tracking data is not available for all individuals.

| Cluster index | Mean X average deviation (std) | Mean Y average deviation (std) | Number of samples with eye-tracking data |
|---|---|---|---|
| 1 | 80.00 (59.26) | 101.20 (92.83) | 27 |
| 2 | 43.53 (29.53) | 66.07 (79.10) | 19 |
| 3 | 58.66 (49.40) | 71.99 (73.12) | 22 |
| 4 | 65.50 (98.81) | 70.61 (93.59) | 47 |
| 5 | 63.50 (49.51) | 59.79 (78.15) | 21 |
| 6 | 53.15 (31.89) | 81.16 (92.24) | 11 |
